# Nanobody-tethered transposition allows for multifactorial chromatin profiling at single-cell resolution

**DOI:** 10.1101/2022.03.08.483436

**Authors:** Tim Stuart, Stephanie Hao, Bingjie Zhang, Levan Mekerishvili, Dan A Landau, Silas Maniatis, Rahul Satija, Ivan Raimondi

## Abstract

Chromatin states are functionally defined by a complex combination of histone modifications, transcription factor binding, DNA accessibility, and other factors. However, most current single-cell-resolution methods are unable to measure more than one aspect of chromatin state in a single experiment, limiting our ability to accurately measure chromatin states. Here, we introduce nanobody-tethered transposition followed by sequencing (NTT-seq), a new assay capable of measuring the genome-wide presence of multiple histone modifications and protein-DNA binding sites at single-cell resolution. NTT-seq utilizes recombinant Tn5 transposase fused to a set of secondary nanobodies (nb). Each nb-Tn5 fusion protein specifically binds to different immunoglobulin-G antibodies, enabling a mixture of primary antibodies binding different epitopes to be used in a single experiment. We apply bulk- and single-cell NTT-seq to generate high-resolution multimodal maps of chromatin states in cell culture and in cells of the human immune system, demonstrating the high accuracy and sensitivity of the method. We further extend NTT-seq to enable simultaneous profiling of cell-surface protein expression alongside multimodal chromatin states to study cells of the immune system.

---

Several related methods were recently developed that enable individual aspects of chromatin state to be measured at single-cell resolution via an antibody-guided DNA tagmentation reaction (Kaya-Okur et al. 2019; Carter et al. 2019; Wang et al. 2019). However, chromatin states are characterized by combinations of factors at an individual locus (Janssen and Lorincz 2021), including histone posttranslational modifications and the binding of non-histone proteins to the DNA. For example, promoters are commonly marked by both H3K27ac and H3K4me2, whereas enhancers are marked by H3K27ac but typically lack H3K4me2. Furthermore, active and poised enhancers are both marked by H3K4me1 and can be distinguished by the presence of H3K27ac (Creyghton et al. 2010). Therefore, multimodal single-cell chromatin profiling methods are required to fully characterize chromatin states in heterogeneous tissues.

A majority of single-cell chromatin profiling methods employ protein-A/G fused to Tn5 transposase (Kaya-Okur et al. 2019; Carter et al. 2019; Wang et al. 2019; Gopalan et al. 2021; Meers et al. 2021). Protein-A/G binds to IgG antibodies, enabling Tn5 to be directed to regions of the genome where an IgG antibody is bound and insert adapters for DNA sequencing. As protein-A/G binds to IgG antibodies from different species with high affinity, such methods are difficult to perform in an antibody-multiplexed design aiming to measure multiple histone modifications in a single experiment. Current approaches for multimodal chromatin profiling using protein-A/G involve complex experimental workflows with multiple wash and incubation steps. These methods have not been demonstrated to work with complex tissues (Gopalan et al. 2021; Meers et al. 2021), thus limiting their broader application. We reasoned that the use of small single polypeptide chain antibodies (nanobodies) that specifically bind IgG from different species or different IgG subtypes in place of protein-A/G may enable the multiplexing of primary antibodies to facilitate a multimodal chromatin assay (Pleiner et al. 2018). Nanobodies bind strongly to their target epitope with dissociation constants (Kd) in the high picomolar-scale, whereas protein-A/G has Kd in the low nanomolar scale (Saha et al. 2003; Hassanzadeh-Ghassabeh et al. 2013). Furthermore, nanobodies are stable under a broad temperature and pH range. We hypothesized that a nanobody-Tn5 (nb-Tn5) fusion would form a stable and specific protein-protein complex with a target primary IgG antibody. Here, we engineered and produced four different recombinant nb-Tn5 fusion proteins, specific for IgG antibodies from different species or IgG subtypes (Fig. 1A, Extended Data Fig. 1A). This included anti-mouse and anti-rabbit IgG nanobodies, as well as isotype-specific nanobodies for mouse IgG1 and IgG2a. Loading nb-Tn5 fusion proteins with barcoded DNA adaptor sequences enables the identity of individual nb-Tn5 fusion proteins that generated the sequenced DNA fragment to be determined through DNA sequencing. We tested each recombinant nb-Tn5 fusion in a bulk-cell NTT-seq experiment and obtained an NTT-seq library only when the nb-Tn5 matched the target antibody, while the incubation of nb-Tn5 with the unmatched Ab resulted in no library amplification via PCR (Extended Data Fig. 1B). Motivated by this result, we performed multiplexed NTT-seq aiming to profile multiple different chromatin features in a single experiment. In our protocol, extracted nuclei are stained in a single step using primary antibodies for multiple epitopes simultaneously, the excess antibody is washed and nuclei are incubated with a mixture of adapter-barcoded (Amini et al. 2014) nb-Tn5s, with each nb-Tn5 recognizing a specific IgG antibody. Subsequently, nb-Tn5s are activated by adding Mg^2+^ resulting in the tagmentation of genomic DNA in proximity of the primary antibody. The released DNA fragments harbor specific barcodes enabling the assignment of sequenced fragments to an individual nb-Tn5 and its associated primary antibody (Fig. 1B).

**Figure 1.**
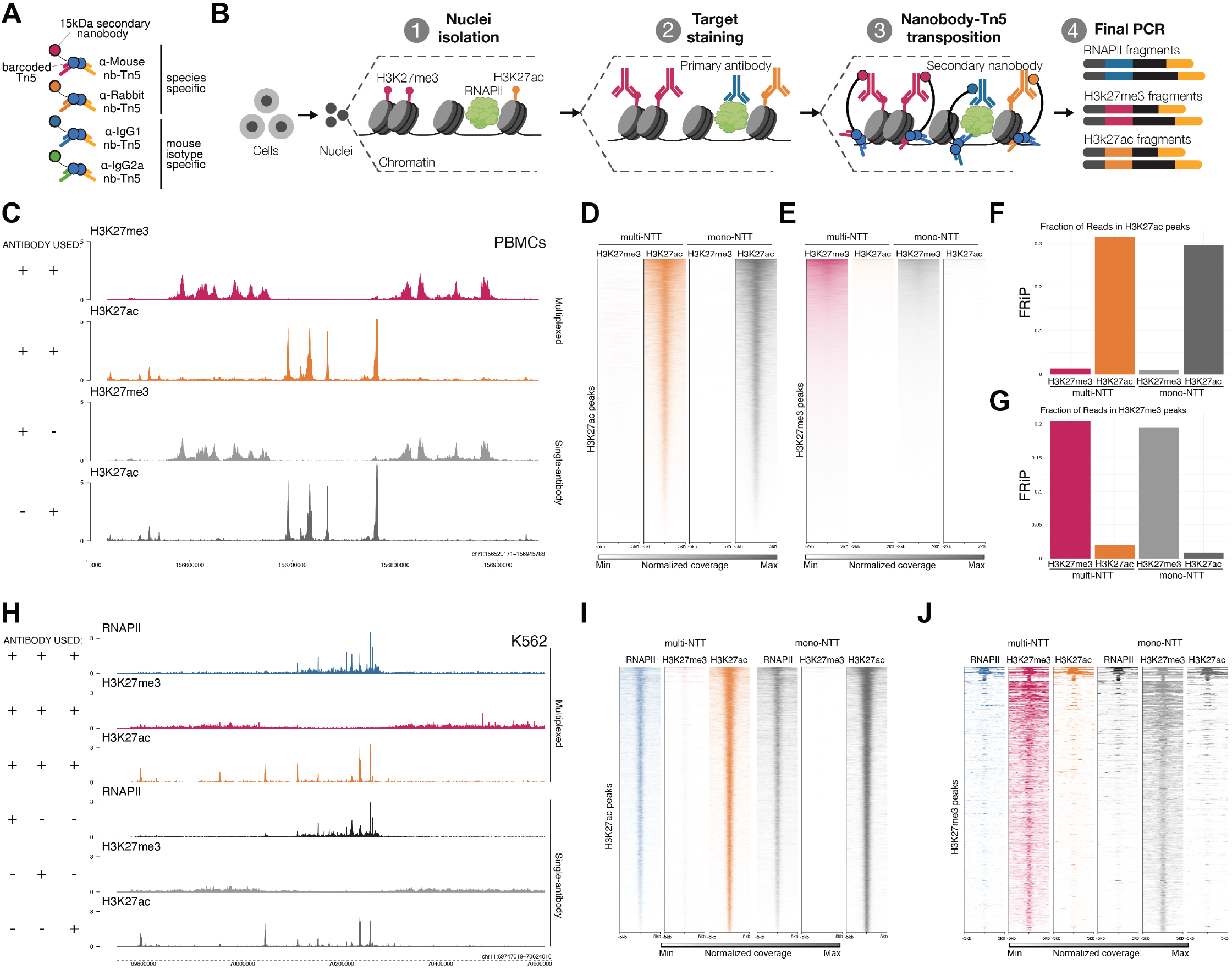
Bulk-cell NTT-seq enables simultaneous profiling of multiple chromatin marks. **A)** Schematic representation of nanobody-Tn5 fusion proteins loaded with barcoded DNA adaptors. **B)** Overview of the NTT-seq protocol. Nuclei are extracted from cells and stained with a mixture of IgG primary antibodies for targets of interest. Nanobody-Tn5 fusion proteins are then added and tagment the genomic DNA surrounding primary antibody binding sites. Released DNA fragments are amplified by PCR to obtain a sequencing library harboring barcode sequences specific for each nb-Tn5 protein used. **C)** Genome browser tracks for a representative region of the human genome. NTT-seq was performed on PBMCs for H3K27me3 alone (light grey), H3K27ac alone (dark grey) or for both together in a multiplexed experiment (red/yellow). Sequencing data were normalized as bins per million mapped reads (BPM) **D)** Heatmap displaying coverage within 33,205 H3K27ac peaks identified using MACS2, for multiplexed (multi) and non-multiplexed (mono) NTT-seq PBMC experiments. **E)** As for D, for 67,459 H3K27me3 peaks. **F)** Fraction of reads in H3K27ac peaks for multiplexed and non-multiplexed NTT-seq PBMC datasets. **G)** As for F, for H3K27me3 peaks. **H)** Genome browser tracks for a representative region of the human genome for multiplexed and non-multiplexed NTT-seq K562 cell datasets. Sequencing data were normalized as bins per million mapped reads (BPM), as for the PBMC datasets. **I)** Heatmap displaying coverage centered on H3K27ac peaks for multiplexed and non-multiplexed NTT-seq experiments using K562 cells, for RNAPII, H3K27ac, and H3K27me3 modalities. **J)** As for I, for H3K27me3 peaks.

To test the targeting specificity of our species-specific nb-Tn5 fusion proteins, we used antibodies for H3K27me3 and H3K27ac in bulk human peripheral blood mononuclear cells (PBMCs), as these marks do not co-occur in the genome (Tie et al. 2009). Multiplexed NTT-seq resulted in libraries with nearly identical genomic distributions for each separate mark to matched NTT-seq performed on the same cells for each histone mark separately (Fig. 1C). The enrichment of sequenced fragments falling in H3K27me3 and H3K27ac peaks was approximately the same across the multiplexed and non-multiplexed experiments (Fig. 1D, E), and showed mutual exclusivity (Fig. 1F, G; Extended Data Fig. 1C). This suggests that multiplexed NTT-seq results in highly accurate localization of chromatin marks genome-wide. Then, we tested our isotype-specific nb-Tn5 profiling of three primary antibodies in a single experiment, repeating similar experiments using K562 cells staining with mouse IgG1 antibody against H3K27me3, mouse IgG2a antibody against H3K27ac, and including an additional rabbit IgG antibody for RNA Polymerase II (RNAPII) with phosphorylated Serine 2 and Serine 5 (elongating RNAPII, enriched on actively transcribed genes) (Zaborowska et al. 2016). In comparison with a control experiment in which each of the three targets was profiled individually, multiplexed NTT-seq again produced comparable specificity of target-specific enrichment in peaks (Fig. 1H, I, J; Extended Data Fig. 1D), demonstrating the ability to profile three targets simultaneously, as well as the ability to profile non-histone proteins.

Encouraged by the results obtained in bulk cells, we next applied NTT-seq to characterize multimodal chromatin states at single-cell resolution using the 10x Genomics scATAC-seq kit (Fig. 2A). We profiled H3K27me3, H3K27ac and elongating RNAPII in a mixture of K562 and HEK293 cells. We obtained on average 743 fragments for H3K27me3, 382 fragments for H3K27ac and 542 fragments for RNAPII per cell, outperforming the recently-developed multiCUT&Tag method (Gopalan et al. 2021) in terms of sensitivity (Extended Data Fig. 2A). We projected cells into a low-dimensional space using latent semantic indexing (LSI) and UMAP (Stuart et al. 2021; Becht et al. 2018), and clustered cells using a weighted combination of all three data modalities (Hao et al. 2021) (Fig. 2B). We identified two groups of cells corresponding to K562 and HEK293 cells. The genomic distribution of reads for each mark obtained in the multiplexed single-cell experiment was highly similar to data from the same cell lines where each feature was profiled individually in bulk (Fig. 2C, Extended Data Fig. 2B). Examining the distribution of fragments at ATAC (ENCODE Project Consortium 2012), H3K27me3, H3K27ac, and RNAPII peaks further showed the co-occupancy of RNAPII and H3K27ac in open chromatin regions, while the signal for H3K27me3 was mutually exclusive with the other profiled marks (Fig. 2D, E). Furthermore, multiplexed single-cell-derived signals were highly correlated with bulk-cell signal for each assay profiled individually (Fig. 2D). Using a combination of cellular modalities provided the strongest separation of the two cell types in low-dimension space. When constructing a neighbor graph, we observed a higher fraction of a cell’s neighbors belonging to the same cell type as that cell when using multiple modalities (Fig. 2F). This highlights the value of multimodal chromatin data in measuring cellular states, and together these results show that NTT-seq is an effective method for profiling multiple chromatin modalities at single-cell resolution.

**Figure 2.**
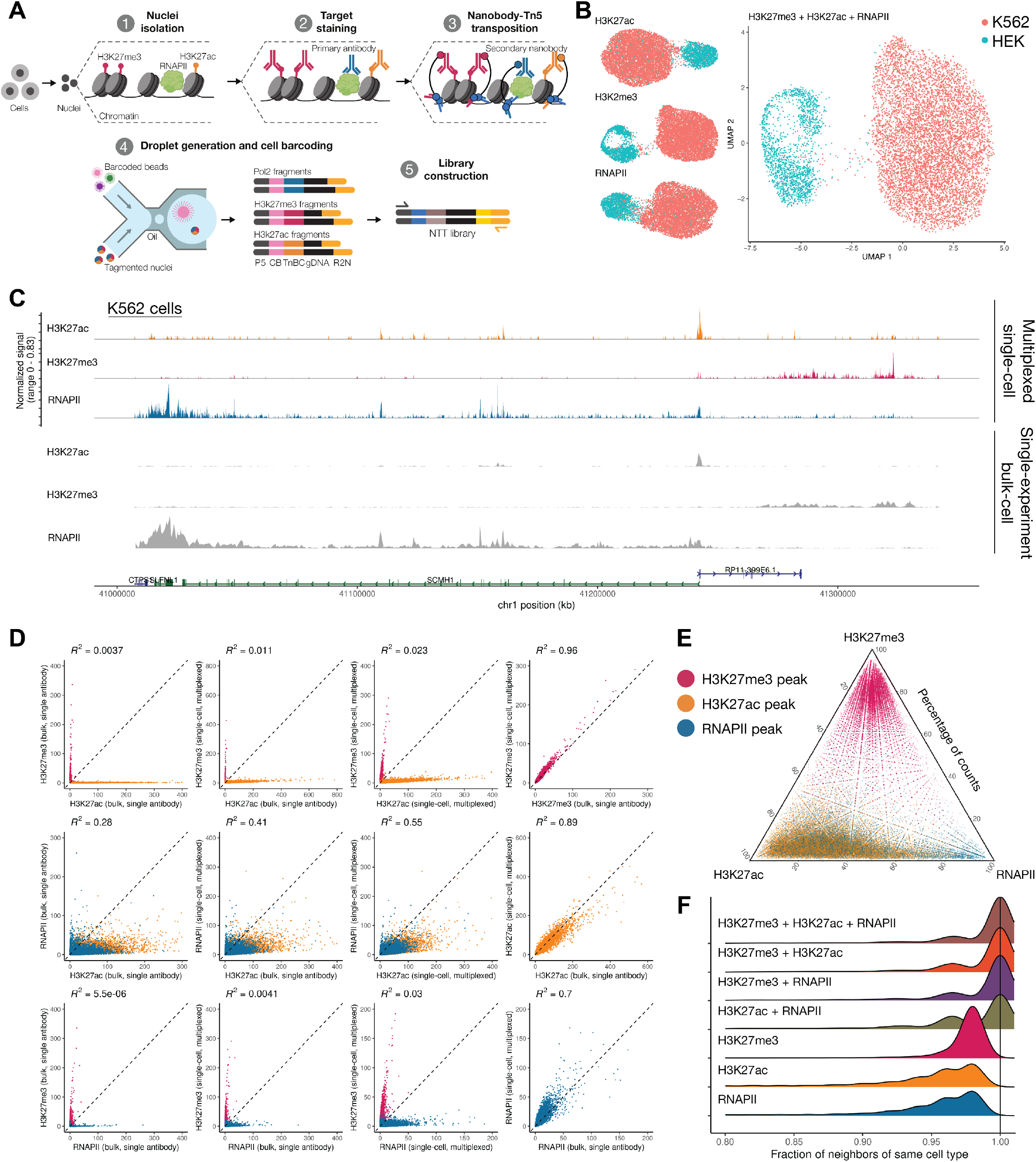
NTT-seq provides accurate single-cell multimodal chromatin profiles. **A)** Schematic overview of the single-cell NTT-seq protocol. Cells are tagmented and processed in bulk (steps 1-3), and are encapsulated in droplets to attach cell-specific barcode sequenced to transposed DNA fragments (steps 4-5). **B)** UMAP representations of cells profiled using multiplexed single-cell NTT-seq. Individual UMAP representations built using each assay are shown (left side), along with a visualization constructed incorporating information from all three chromatin modalities (WNN UMAP, right side). Cells are colored by their predicted cell type. **C)** Multimodal genome browser view of a representative genomic locus, for K562 cells. Top three tracks show H3K27ac, H3K27me3, and RNAPII profiled simultaneously in a single-cell experiment. Lower three tracks show H3K27ac, H3K27me3, and RNAPII profiled individually in bulk-cell NTT-seq experiments using K562 cells. **D)** Scatterplots showing normalized fragment counts for H3K27me3, H3K27ac, and RNAPII peaks defined by ENCODE (ENCODE Project Consortium 2012), for bulk and single-cell multiplexed NTT-seq experiments, for K562 cells. Peaks are colored according to their chromatin modality (red: H3K27me3 peak, yellow: H3K27ac peak, blue: RNAPII peak). Coefficient of determination (*R*2) between experiments are shown above each scatterplot. **E)** Ternary plot showing the relative frequency of H3K27me3, H3K27ac, and RNAPII fragment counts within H3K27me3, H3K27ac, and RNAPII peak regions defined by ENCODE ChIP-seq datasets. **F)** Fraction a cell’s nearest-neighbors belonging to the same predicted cell type, for neighbor graphs defined using a single chromatin modality or a weighted combination of modalities.

We next sought to extend the NTT-seq method to enable simultaneous measurement of cell surface protein expression alongside multimodal chromatin states at single-cell resolution. Building on the recently-developed CUT&Tag-pro method (Zhang et al. 2021), we stained a population of mobilized PBMCs with an oligonucleotide-conjugated panel of 173 antibodies targeting immune-relevant cell surface proteins. Cells were then crosslinked, permeabilized, and incubated with antibodies against H3K27me3 and H3K27ac, and our standard NTT-seq protocol followed to generate single-cell libraries. This resulted in a dataset of 4,870 cells with a mean of 2,732 H3K27me3 and 398 H3K27ac fragments per cell (Extended Data Fig. 3A), as well 666 antibody-derived tag (ADT) counts per cell, similar to sensitivity recently demonstrated for cell surface protein quantified alongside a single chromatin mark (Extended Data Fig. 3A) (Zhang et al. 2021). We clustered cells using a weighted combination of each modality (Hao et al. 2021) and annotated cell clusters based on their patterns of protein expression (Fig. 3A). Protein expression patterns were concordant with cell clusters determined from a chromatin-based clustering, and we observed uniform expression of CD3 in T cells, mutually exclusive expression of CD4 and CD8, expression of CD14 in monocytes, CD19 in B cells, and IL2RB in NK cells (Fig. 3B). Pseudobulk H3K27me3 and H3K27ac NTT-seq profiles were highly correlated with individual single-cell CUT&Tag-pro (Zhang et al. 2021) profiles for human PBMCs for the same histone marks (Fig. 3C). Consistent with our previous results, we also observed an extremely low coefficient of determination (*R*^2^=0.00028) between H3K27me3 and H3K27ac levels within peaks (Fig. 3D), further supporting the accuracy of multiplexed NTT-seq single-cell profiles when applied to complex tissues. We further observed consistency between chromatin states and protein expression patterns for each cell type, supporting accurate cell-surface protein quantification. For example, the *PAX5* locus was repressed in non-B cells with low CD19 protein expression, and active in B cells with high CD19 expression (Fig. 3E). Similarly, the *CD33* locus was active in monocytes with high CD33 protein expression and repressed in B cells with low CD33 expression. To evaluate the accuracy of our cell type classifications and multimodal chromatin landscapes measured by NTT-seq, we compared the results of our single-cell NTT-seq experiment with FACS-sorted ChIP-seq profiles for CD14 monocytes, HSCs, and B cells previously published by the ENCODE consortium (ENCODE Project Consortium 2012). Pseudobulk profiles generated from our NTT-seq cell types recapitulated the expected cell-type-specific ENCODE ChIP-seq profiles (Extended Data Fig. 3B).

**Figure 3.**
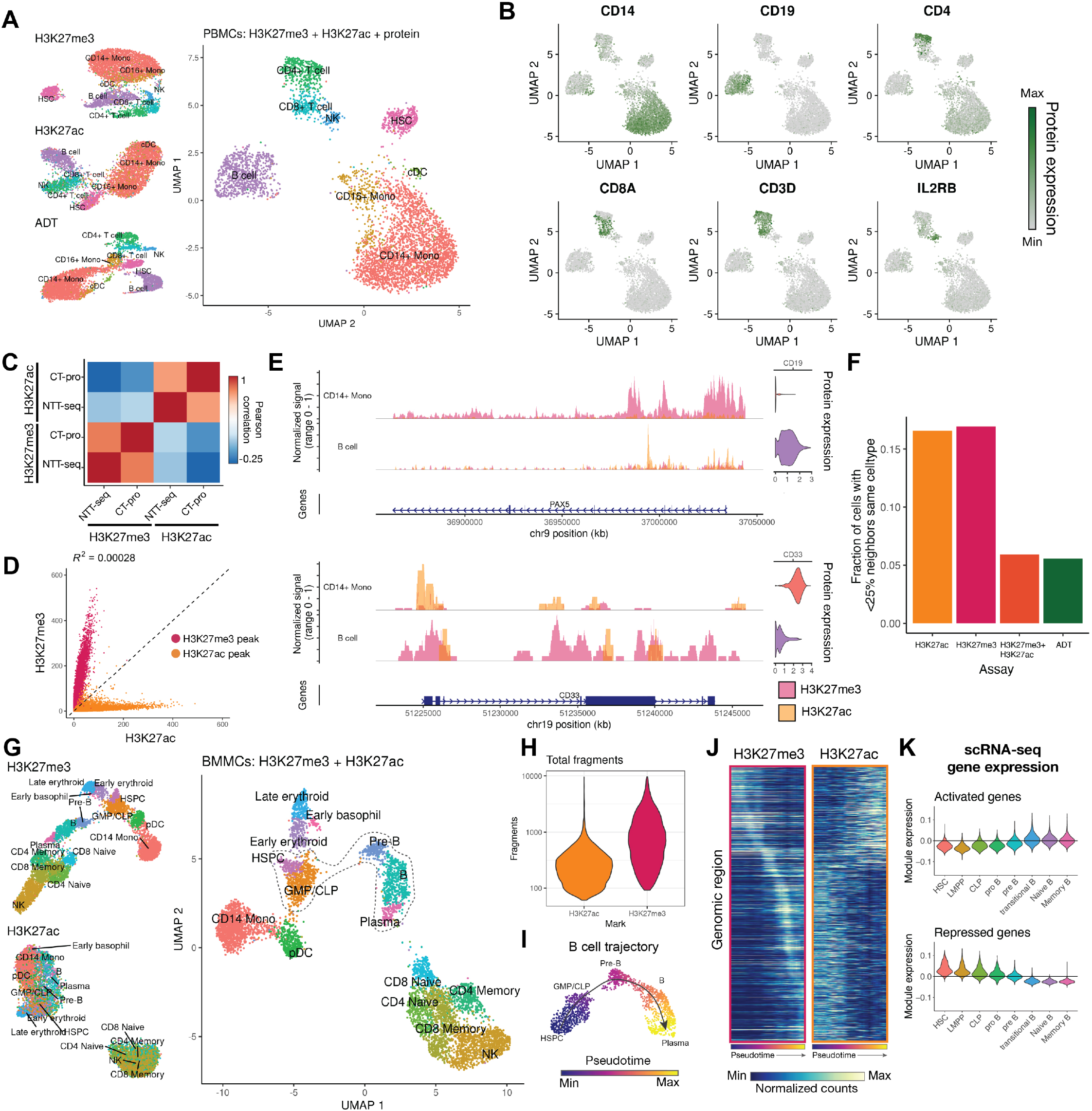
Application of multiplexed single-cell NTT-seq to human tissues. **A)** UMAP representation of PBMCs profiled using NTT-seq with protein expression. Separate UMAP representations constructed separately for each assay are shown (left side), along with a multimodal UMAP constructed using all three data modalities (right side). Cells are colored and labeled by their annotated cell types. **B)** Patterns of cell-surface-protein expression in human PBMCs profiled using NTT-seq. **C)** Pearson correlation between genome-wide NTT-seq and scCUT&Tag-pro (CT-pro) signal in PBMCs within H3K27me3 and H3K27ac peaks. **D)** Scatterplot showing the number of counts per H3K27me3 and H3K27ac peak for each assay, for PBMCs profiled using single-cell multiplexed NTT-seq. Peaks are colored according to their assay (red: H3K27me3 peaks; yellow: H3K27ac peaks). Coefficient of determination (*R*2) is shown above the plot. Axes show total fragment counts per million within individual peaks. **E)** Multimodal genome browser view of the *PAX5* and *CD33* loci, for B cells and CD14+ monocytes. Normalized protein expression values are shown alongside coverage tracks for each cell type, for CD19 and CD33 protein. H3K27me3 and H3K27ac histone modification profiles are overlaid with transparency, with the signal for each scaled to the maximal signal within the genomic region shown. **F)** Fraction of cell with <25% of neighbors belonging to the same cell type, for neighbor graphs defined using individual chromatin modalities, cell-surface protein expression, or a combination of chromatin modalities. **G)** UMAP representation of BMMCs profiled using multiplexed single-cell NTT-seq. Separate UMAP representations constructed for H3K27me3 and H3K27ac separately are shown (left side), and a UMAP representation constructed using a weighted combination of H3K27me3 and H3K27ac measurements is shown (right). Cells are colored and labeled by their annotated cell type. HSPC: hematopoietic stem and progenitor cells; GMP/CLP: granulocyte monocyte progenitor / common lymphoid progenitor; CD14 Mono: CD14+ monocyte; pDC: plasmacytoid dendritic cell; NK: natural killer cell. **H)** Distribution of total fragment counts per cell for H3K27ac and H3K27me3. Counts are displayed using a log10 scale. **I)** Pseudotime trajectory for B cell development. HSPCs, GMP/CLP, pre-B, B, and plasma cells were subset from the main dataset and used to construct a new UMAP visualization containing only these cells. Cells are colored by their pseudotime value and labeled by their annotated cell type. **J)** Heatmap showing H3K27me3 and H3K27ac signal for 10 kb genomic bins correlated with B cell pseudotime progression. Heatmaps show the same genomic regions for both assays, with identical ordering of genomic regions. **K)** Expression of genes close to activated (gain H3K27ac, upper plot) or repressed (gain H3K27me3, lower plot) genomic regions in a separate scRNA-seq BMMC dataset, for cells in the B cell developmental trajectory.

While cell-surface protein expression information provides a powerful method of studying immune cells, these methods are of limited value outside of the immunology field. To test whether a low-dimensional structure similar to that obtained using protein expression could be learned using the chromatin data alone, we compared the neighbor graphs obtained using protein expression data to that obtained using individual or combined chromatin modalities. While individual chromatin marks were unable to faithfully recapitulate the low-dimensional structure observed when including protein expression data, the combination of H3K27me3 and H3K27ac modalities provided a similar low-dimensional neighbor structure (Fig. 3F). This again highlights the unique power of multimodal chromatin data in resolving cellular states, and indicates that multiplexed NTT-seq may be a powerful method capable of characterizing heterogeneous tissues without the need for cell surface protein measurements.

We next sought to apply NTT-seq in a complex tissue that contains differentiating cells to capture chromatin remodeling dynamics that shape cellular identity. We profiled H3K27me3 and H3K27ac in human bone marrow mononuclear cells (BMMCs) (Fig. 3G). This resulted in 6,192 cells with a mean of 1,084 and 292 fragments per cell for H3K27me3 and H3K27ac respectively (Fig. 3H). We annotated cell clusters using a combination of label transfer using an annotated BMMC scATAC-seq dataset (Granja et al. 2019; Stuart et al. 2019) using the H3K27ac assay, and manual annotation inspecting the presence of active and repressive histone marks at key marker genes for each cell type. We identified the expected cell types present in the immune system, including hematopoietic stem and progenitor cells (HSPCs) (Fig. 3G). Consistent with results obtained using cells in culture and PBMCs, we observed mutual exclusivity between H3K27ac and H3K27me3 across regions of the genome for BMMCs (Extended Data Fig. 3C). To study how multimodal chromatin states may change during cell development, we ordered cells belonging to the B cell lineage, including HSPCs, common lymphoid progenitors (CLPs), pre-B, B, and plasma cells along a developmental pseudotime trajectory using Monocle 3 (Qiu et al. 2017) (Fig. 3I). This revealed the expected ordering of cells in a trajectory leading from HSPCs through CLP, pre-B, B, and plasma cells. To identify regions of the genome that changed their H3K27me3 and H3K27ac state across this trajectory, we quantified fragment counts for each cell in 10 kb bins spanning the entire genome for each chromatin modality. We identified genome bins with signal correlated with pseudotime, and identified a set of 988 regions with opposing relationships between H3K27me3 and H3K27ac signal (i.e., a gain in signal for one assay was accompanied by a loss of signal for the other assay). Sorting these regions by the point at which they reached maximal H3K27me3 signal revealed an ordered sequence of sites that became repressed or activated during B cell development (Fig. 3J). The genome bin with the strongest opposing H3K27ac and H3K27me3 signals across pseudotime was located at the *PAX5* promoter, a B-cell-specific transcription factor. To systematically assess the cell-type-specific expression pattern of genes located near genomic bins that were repressed or activated along the B cell pseudotime trajectory, we examined a published scRNA-seq dataset for healthy human BMMCs. We identified the closest gene to each pseudotime-correlated genome bin, and classified these as activated (positive correlation between H3K27ac and pseudotime) or repressed (positive correlation between H3K27me3 and pseudotime). Examining the expression of repressed and activated genes in the scRNA-seq dataset revealed concordant patterns of gene expression, with chromatin-activated genes becoming expressed later in B cell development, and repressed genes being expressed in HSPCs but turned off later in B cell development (Fig. 3K).

Together these analyses demonstrate that NTT-seq datasets provide accurate multimodal chromatin landscapes at single-cell resolution, contain sufficient information to identify major cell types and states in complex human tissues, provide profiles that reflect high-quality bulk ChIP-seq data (ENCODE Project Consortium 2012), and can be generated in conjunction with accurate cell-surface protein expression measurements. Existing multimodal chromatin technologies require complex experimental workflows and have not been demonstrated to work with complex tissue samples (Gopalan et al. 2021; Meers et al. 2021), or are strictly limited in the chromatin states that they can measure (Tedesco et al. 2021). NTT-seq overcomes both of these limitations, providing a streamlined experimental workflow applicable to complex tissues, and is limited only by the availability of primary antibodies. We anticipate future reagent development and protocol improvements will enable us to scale NTT-seq to profiling of more than three markers simultaneously, and are actively working on the generation of additional nb-Tn5s targeting antibodies raised in different species such as goat, rat, sheep, guinea pig and multiple IgG isotypes within the same species. This will expand the portfolio of reagents for multimodal chromatin profiling. Moreover, we anticipate that the use of dual-barcoded nb-Tn5 can be implemented in our protocol to investigate intra-locus interactions between different chromatin features, such as bivalent promoters or enhancers. We believe that the simplicity with which NTT-seq achieves simultaneous profiling of chromatin features makes this approach particularly appealing, and could represent the standard for multifactorial chromatin mapping in the future.

## Code availability

Signac 1.6.0 (Stuart et al. 2021) and Seurat 4.1.0 (Hao et al. 2021) was used for all analysis, and are available from CRAN: https://cran.r-project.org/package=Signac; https://cran.r-project.org/package=Seurat. Code to reproduce analyses is available on GitHub: https://github.com/timoast/nanobody.

## Methods

### Cell culture

K562 cells were acquired from ATCC (nos. CCL-243). HEK293FT cells were acquired from Thermo Fisher (no. R70007). HEK293FT cells were maintained at 37°C and 5% CO_2_ in D10 medium (DMEM with high glucose and stabilized L-glutamine (Caisson, no. DML23) supplemented with 10% fetal bovine serum (FBS; Thermo Fisher, no. 16000044)). K562 cells were maintained at 37°C and 5% CO_2_ in R10 medium (RPMI with stabilized L-glutamine (Thermo Fisher, no. 11875119) supplemented with 10% FBS).

### Primary cells acquisition and processing

Cryopreserved healthy donor PBMCs were isolated from mobilized peripheral blood, BMMCs were purchased from AllCells. After thawing into DMEM with 10% FBS, the cells were spun down at 4°C for 5 min at 400 g and washed twice with PBS with 2% BSA. After centrifugation, the cell pellet was resuspended in staining buffer (2% BSA and 0.01% Tween in PBS).

### Cloning of nb-Tn5 plasmid constructs

Previously published sequences coding for secondary nanobodies (Pleiner et al. 2018) were synthesized as a gene fragment (IDT) flanked by restriction enzyme sites NcoI and EcoRI. To replace protein-A with a nanobody, 3XFlag-pA-Tn5-Fl (addgene #124601) and gene fragments were digested with NcoI and EcoRI 1h at 37°C, ligated overnight at 16°C and subsequently transformed into competent cells (NEB C2992H).

### Nanobody-Tn5 transposase production

The pTXB1-nbTn5 vector was transformed into BL21(DE3)-competent *Escherichia coli* cells (NEB, no. C2527), and nb-Tn5 was produced via intein purification with an affinity chitin-binding tag (Picelli et al. 2014). 400 mL of Luria broth (LB) culture was grown at 37°C to optical density (OD600)=0.6. nb-Tn5 expression was then induced with isopropyl-ß-d-thiogalactopyranoside (IPTG) 0.25mM at 22°C 6 hours. After induction, cells were pelleted and then frozen at −80°C overnight. Cells were then lysed by sonication in 100 mL pf HEGX (20mM HEPES-KOH pH 7.5, 0.8M NaCl, 1mM EDTA, 10% glycerol, 0.2% Triton X-100) with a protease inhibitor cocktail (Roche, no. 04693132001). The lysate was pelleted at 30,000g for 20 min at 4°C. The supernatant was transferred to a new tube, and 3 µL of neutralized 8.5% polyethylenimine (Sigma-Aldrich, P3143) was added dropwise to each 100 µL of bacterial extract, gently mixed and centrifuged at 30,000g for 30 min at 4°C to precipitate DNA. The supernatant was loaded on four 2 mL chitin columns (NEB, no. S6651S). Columns were washed with 10 mL of HEGX, then 1.5 mL of HEGX containing 100 mM DTT was added to the column with incubation for 48h at 4°C to allow cleavage of nb-Tn5 from the intein tag. nb-Tn5 was eluted directly into two 30 kDa molecular-weight cutoff (MWCO) spin columns (Millipore, no. UFC903008) by the addition of 2 mL of HEGX. Protein was dialyzed in five dialysis steps using 15 mL of 2x dialysis buffer (100 HEPES-KOH pH 7.2, 0.2M NaCl, 0.2 mM EDTA, 2 mM DTT, 20% glycerol) and concentrated to 1 mL by centrifugation at 5,000g. The protein concentrate was transferred to a new tube and mixed with an equal volume of 100% glycerol. nb-Tn5 aliquots were stored at −80°C.

### Transposome assembly

We obtained barcoded Tn5 adaptors from IDT, as described by Amini et al. (2014) with 8 bp barcodes designed using FreeBarcodes (Hawkins et al., 2018). To produce mosaic-end, double-stranded (MEDS) oligos, we annealed each barcoded T5 tagmentation oligo with the pMENT common oligo (100µM each) as follows, in TE buffer: 95°C for 5min then cooling at 0.2°Cs–1 to 4°C (bcMEDS-A). The same process was used to anneal single T7 tagment oligo with the pMENT common oligo (MEDS B; Supplementary Table). bcMEDS-A and MEDS-B were mixed 1:1, 6µL was transferred to a new tube and mixed with 10µL of nb-Tn5 enzyme. After 1 hour at room temperature to allow for transposome assembly.

### Antibodies

Antibodies used were H3K27ac (1:50, Active Motif, 39133), H3K27ac (1:50, Active Motif, 91193), H3K27me3 (1:50, Active Motif, 61017), Phospho-Rpb1 CTD (Ser2/Ser5) (1:50, Cell Signaling, 13546). For NTT-seq with surface markers readout on primary cells, TotalSeq-A conjugated panel were obtained from BioLegend (399907).

## NTT-seq

### Antibody staining

For NTT-seq with surface markers readout on primary cells, 1 million thawed PBMC or BMMC were resuspended in 200 µL staining buffer (2% BSA and 0.01% Tween in PBS) and incubated for 15 min with 20 µL Fc receptor block (TruStain FcX, BioLegend) on ice. Cells were then washed three times with 1 mL staining buffer and pooled together. The panel of oligo-conjguated antibodies was added to the cells to incubate for 30 min on ice. After staining, cells were washed three times with 1 mL staining buffer and resuspended in 100 µL staining buffer. After the final wash, cells were resuspended 200 µL PBS ready for fixation.

### Fixation and permeabilization

For human cell lines, nuclei were extracted as previously described (Kaya-Okur et al. 2020) and resuspended in 150 µL of PBS. Then, 16% methanol-free formaldehyde (Thermo Fisher Scientific, PI28906) was added for fixation (final concentration: 0.1%) at room temperature for 3 min. The cross-linking reaction was stopped by addition of 12 µL 1.25 M glycine solution. Subsequently, nuclei were washed once with 150 µL antibody buffer (20 mM HEPES pH 7.6, 150 mM NaCl, 2 mM EDTA, 0.5 mM spermidine, 1% BSA, 1× protease inhibitors).

For NTT-seq with surface markers readout on primary cells, 16% methanol-free formaldehyde (Thermo Fisher Scientific, PI28906) was added for fixation (final concentration: 0.1%) at room temperature for 5 min. The cross-linking reaction was stopped by addition of 12 µL 1.25 M glycine solution. Subsequently, cells were washed twice with PBS. The permeabilization was performed by adding isotonic lysis buffer (20 mM Tris-HCl pH 7.4, 150 mM NaCl, 3 mM MgCl_2_, 0.1% NP40, 0.1% Tween-20, 1% BSA, 1× protease inhibitors) on ice for 7 min. Subsequently, 1 mL of cold wash buffer (20 mM HEPES pH 7.6, 150 mM NaCl, 0.5 mM spermidine, 1× protease inhibitors) was added, and cells were centrifuged at 800g for 5 min at 4°C.

### Tagmentation

Nuclei or permeabilized cells were directly suspended with 150 µL antibody buffer (20 mM HEPES pH 7.6, 150 mM NaCl, 2 mM EDTA, 0.5 mM spermidine, 1% BSA, 1× protease inhibitors) with a cocktail of primary antibodies and incubated overnight on a rotator at 4°C. The next day cells were washed twice with 150 μL wash buffer to remove the remaining antibodies. The cells were then resuspended in 150 μL high salt wash buffer (20 mM HEPES pH 7.6, 300 mM NaCl, 0.5 mM spermidine, 1× protease inhibitors) with 2.5 µL nb-Tn5 for each target of interest and incubated for 1 h on a rotator at room temperature. The cells were then washed twice with high salt wash buffer and resuspended in 50 μL tagmentation buffer (20 mM HEPES pH 7.6, 300 mM NaCl, 0.5 mM spermidine, 10 mM MgCl2, 1× protease inhibitors). The samples were incubated for 1 h at 37°C. Tagmentation steps were performed in 0.2 mL tubes to minimize cell loss.

### Bulk-cell NTT-seq

To stop tagmentation, 1µL of 0.5M EDTA, 1µL of 10% SDS and 0.25µL of 20mg/mL Proteinase K was added to the sample, incubated at 55°C for 1 hour. DNA was extracted with Chip DNA clean & Concentrator kit (Zymo Research, D5201) following manufacturer instructions. To amplify libraries, 21µL DNA was mixed with 2µL of a universal i5 and a uniquely barcoded i7 primer, using a different barcode for each sample. A volume of 25µL NEBNext HiFi 2× PCR Master mix was added and mixed. The sample was placed in a Thermocycler with a heated lid using the following cycling conditions: 72°C for 5min (gap filling); 98°C for 30s; 14 cycles of 98°C for 10s and 63°C for 30s; final extension at 72°C for 1min and hold at 8°C. Post-PCR clean-up was performed by adding 1.1× volume of Ampure XP beads (Beckman Coulter), and libraries were incubated with beads for 15min at RT, washed twice gently in 80% ethanol, and eluted in 30µL 10mM Tris pH 8.0.

### NTT-seq single cell encapsulation, PCR, and library construction

After tagmentation, cells were centrifuged for 5min at 1,000g and the supernatant was discarded. Cells were resuspended with 30 µL 1× Diluted Nuclei Buffer (10x Genomics, #2000207), counted, and diluted to a concentration based on the targeted cell number. The transposed cell mix was prepared as following: 7µL of ATAC buffer and 8 µL cells in 1× Diluted Nuclei Buffer. All remaining steps were performed according to the 10x Chromium Single Cell ATAC protocol. For NTT-seq with surface markers readout on primary cells, the library construction method was adapted from ASAP-seq (Mimitou et al. 2021). Briefly, 0.5 μL of 1 μM bridge oligo A (TCGTCGGCAGCGTCAGATGTGTATAAGAGACAGNNNNN NNNNVTTTTTTTTTTTTTTTTTTTTTTTTTTTTTT/3InvdT/) was added to the barcoding mix. Linear amplification was performing using the following PCR program: (40°C for 5 min, 72°C for 5 min, 98°C for 30 s; 12 cycles of 98°C for 10 s, 59°C for 30 s and 72°C for 1 min; ending with hold at 15°C). The remaining steps were performed according to the 10x scATAC-seq protocol (v1.1), with the following additional modifications:

Antibody-derived tags: during silane bead elution (Step 3.1s), beads were eluted in 43.5 μL of elution solution I. The extra 3 μL was used for the surface protein tags library. During SPRI cleanup (Step 3.2d), the supernatant was saved and the short DNA derived from antibody oligos was purified with 2× SPRI beads. The eluted DNA was combined with the 3 µL left aside after the silane purification to be used as input for protein tag amplification. PCR reactions were set up to generate the protein tag library with Kapa Hifi Master Mix (P5 and RPI-x primers): 95°C for 3 min; 14–16 cycles of 95°C for 20 s, 60°C for 30 s and 72°C for 20 s; followed by 72°C for 5 min and ending with hold at 4°C.

RPI-x primer: CAAGCAGAAGACGGCATACGAGATxxxxxxxxGTGACTGG AGTTCCTTGGCACCCGAGAATTCCA.

P5 Primer: AATGATACGGCGACCACCGAGATCTACAC

### Sequencing

The final libraries were sequenced on NextSeq 550 by using custom primers (Amini et al. 2014) with the following strategy: i5: 38bp, i7: 8bp, read1: 60bp, read2: 60bp.

### Bulk-cell data analysis

Bulk-cell data for the cell culture and PBMC datasets were mapped to the hg38 analysis set using Chromap (Zhang et al. 2021), with the –preset atac option set. Output BED files produced by Chromap were sorted, bgzip-compressed, and indexed using tabix for further analysis. Peaks were called using MACS2 (Zhang et al. 2008), with the following options: –nomodel –shift -100 –extsize 200 –format BED; for H3K27me3 data the –broad option was also set. We created bigWig files for each experiment by first creating a bedgraph file using the bedtools genomecov function (Quinlan and Hall 2010) followed by the UCSC bedGraphToBigWig function. Multi-NTT-seq heatmaps were generated in DeepTools (Ramirez et al. 2014).

ChIP-seq peak coordinates for H3K27me3 and H3K27ac for bulk PBMCs, and for H3K27me3, H3K27ac, and RNAPII serine-2 and serine-5 phosphate for K562 cells were downloaded from ENCODE (ENCODE Project Consortium 2012). We counted sequenced DNA fragments falling within each peak region for each bulk-cell PBMC or K562-cell NTT-seq dataset using custom R code and the scanTabix function in Rsamtools, and normalized counts according to the total number of mapped reads for each dataset (counts per million mapped reads normalization). The coefficient of determination (*R*^2^) between peak counts across pairs of experiments was computed using the lm function in R.

## Single-cell data analysis

### Cell culture dataset

#### Read mapping

Reads were mapped to the hg38 analysis set using bwa-mem2 (Vasimuddin et al. 2019) with default parameters, the output sorted and indexed using samtools (Li et al. 2009), and the resulting BAM file used to create a fragment file using the Sinto package (https://github.com/timoast/sinto). We ran the sinto fragmvvents command with the -- barcode_regex “[^:]*” parameter set to extract cell barcodes from the read name. Output files were coordinate-sorted, bgzip-compressed and indexed using tabix (Li 2011), and the resulting fragment files used as input to downstream analyses.

#### Quantification, quality control, and dimension reduction

Genomic regions were quantified using the AggregateTiles function in Signac (Stuart et al. 2021) with binsize=10000 and min_counts=1, using the hg38 genome. Cells with <10,000 total counts, >75 H3K27ac counts, >150 H3K27me3 counts, and >100 RNAPII counts were retained for further analysis. Each assay was processed by performing TF-IDF normalization on the count matrix for the assay, followed by latent semantic indexing (LSI) using the RunTFIDF and RunSVD functions in Signac with default parameters. Two-dimensional visualizations were created for each assay using UMAP, using LSI dimensions 2 to 10 for each assay. Weighted nearest neighbor (WNN) analysis was performed using the FindMultiModalNeighbors function in Seurat, with reduction.list = list(“lsi.k27ac”, “lsi.k27me”, “lsi.pol2”) and dims = list(2:10, 2:10, 2:10) to use LSI dimensions 2 to 10 for each assay. Cell clustering was performed using the resulting WNN graph using the Smart Local Moving community detection algorithm (Waltman and van Eck 2013) by running the FindClusters function in Seurat, with algorithm=3, graph.name=“wsnn”, and resolution=0.05. This resulted in two cell clusters, which were assigned as HEK or K562 based on their correlation with bulk-cell chromatin data for HEK and K562 cells.

#### Specificity analysis

K562-cell bulk ChIP-seq peaks for H3K27ac, H3K27me3, and RNA Pol2 Ser-2 and Ser-5 phosphate were downloaded from ENCODE (ENCODE Project Consortium 2012). Ser-2 and Ser-5 phosphate peaks were combined using the reduce function from the GenomicRanges R package. Fragment counts for K562 cells in the bulk and single-cell dataset were quantified for each peak using the scanTabix function in the Rsamtools R package, with counts normalized according to the total sequencing depth for each dataset. To assess the targeting specificity in single-cell NTT-seq, we computed the coefficient of determination (*R*^2^) between peak counts for each pair of assays, and between bulk and single-cell data for the same assay. We visualized relative peak counts for each assay for each peak by creating a ternary plot using the ggtern R package (Hamilton and Ferry 2018). To assess the low-dimensional neighbor structure obtained using each assay or combinations of assays, we computed the fraction of *k*-nearest neighbors for each cell *i* that belonged to the same cell type classification as cell *i* (*k*=50 for single-modality neighborhoods, variable *k* per-cell for multimodal neighbor graph due to the weighted nearest neighbor method).

#### multi-CUT&Tag comparison

To create a fragment file for the published multi-CUT&Tag dataset, raw sequencing data from Gopalan et al. (2021) were downloaded from NCBI SRA and split into separate FASTQ files according to their Tn5 barcode using a custom Python script. Reads were mapped to the hg38 genome using bwa-mem2 and fragment files created as described above for the NTT-seq datasets. Code to reproduce this analysis is available on GitHub: https://github.com/timoast/multi-ct. We ran the CountFragments function in Signac to count the total number of fragments per cell for each multi-CUT&Tag assay, and retained cells with >200 total counts for further analysis, as described in the original publication (Gopalan et al. 2021). For mixed-barcode fragments we counted ½ count to the total of each assay matching the pair of Tn5 barcodes.

### PBMC dataset

#### Read mapping

Raw genomic reads were mapped and processed as described above for the cell culture single-cell dataset. Antibody-derived tag (ADT) reads were processed using Alevin (Srivastava et al. 2019). We first created a salmon index (Patro et al. 2017) for the BioLegend TotalSeq-A antibody panel, with the --features -k7 parameters. We quantified counts for each ADT barcode using the salmon alevin command with the following parameters: -- naiveEqclass, --keepCBFraction 0.8, --bc-geometry 1[1-16], --umi-geometry 2[1-10], --read-geometry 2[71-85].

#### Quantification, quality control, and dimension reduction

Genomic bins were quantified using the AggregateTiles function in Signac, with binsize=5000 and min_counts=1 to quantify 5 kb bins genome-wide, retaining bins with at least one count. We retained cells with <40,000 and >300 H3K27me3 counts, <10,000 and >100 H3K27ac counts, and <10,000 and >100 antibody-derived tag (ADT) counts. We normalized the ADT data using a centered log ratio transformation using the NormalizeData function in Seurat, with normalization.method=“CLR” and margin=2. We reduced the dimensionality of the ADT assay by first scaling and centering the protein expression values, and running PCA (ScaleData and RunPCA functions in Seurat). We computed a 2-dimensional UMAP visualization using the first 40 principal components (PCs), and clustered cells using the Louvain community detection algorithm. We identified and removed two low-quality clusters containing higher overall ADT counts, as well as higher counts for naive IgG antibodies included in the staining panel. After removing low-quality ADT clusters, reduced the dimensionality of the H3K27me3 and H3K27ac assays using LSI (FindTopFeatures, RunTFIDF, RunSVD functions in Signac) and UMAP using LSI dimensions 2 to 30 for each chromatin assay. To construct a low-dimensional representation using all three data modalities, we ran the weighted nearest neighbors (WNN) algorithm, using the first 40 ADT PCs, and LSI dimensions 2 to 30 for H3K27me3 and H3K27ac (FindMultiModalNeighbors function in Seurat). We clustered cells using the WNN neighbor graph using the Smart Local Moving algorithm (Waltman and van Eck 2013) (FindClusters function in Seurat with algorithm=3 and resolution=1). Cell clusters were manually annotated cell types using the protein expression information. To compare the low-dimensional structure obtained using individual chromatin modalities or combinations of modalities, we computed for each cell *i* the fraction of neighboring cells annotated as the same cell type as cell *i*. We repeated this computation using neighbor graphs computed using single data modalities, or weighted combinations of modalities computed using the WNN method.

#### ENCODE data comparison

Peaks and genomic coverage bigWig files for H3K27me3 and H3K27ac ChIP-seq published by the ENCODE consortium (ENCODE Project Consortium 2012) for B cells, CD34+ CMPs, and CD14+ monocytes were downloaded from the ENCODE website (https://www.encodeproject.org/). We created bigWig files for each corresponding cell type identified in the single-cell multiplexed NTT-seq PBMC dataset by writing sequenced fragments for those cells to a separate BED file, creating a bedGraph file using the bedtools genomecov command (Quinlan and Hall 2010), and creating a bigWig file using the UCSC bedGraphToBigWig tool. We computed the genomic coverage for NTT-seq datasets and ChIP-seq datasets within H3K27me3 and H3K27ac regions using the deeptools multiBigwigSummary function (Ramirez et al. 2014) with the –outRawCounts option set to output the raw correlation matrix as a text file. We computed the correlation between peak region coverage in NTT-seq and ENCODE ChIP-seq datasets using the cor function in R with method=“spearman”.

#### CUT&Tag-pro data comparison

Processed CUT&Tag-pro H3K27me3 and H3K27ac datasets for human PBMCs were downloaded from Zenodo: https://zenodo.org/record/5504061. We compared the number of antibody-derived tag (ADT) counts in NTT-seq and scCUT&Tag-pro datasets by extracting the total number of ADT counts per cell from the scCUT&Tag-pro and NTT-seq Seurat objects and plotting the distribution of total ADT counts per cell for each dataset. We created bigWig files for each scCUT&Tag-pro dataset by first creating a bedGraph file using the bedtools genomecov function, and then creating a bigWig file using the UCSC bedGraphToBigWig function. We computed the coverage for scCUT&Tag-pro datasets within H3K27me3 and H3K27ac PBMC ENCODE peaks using the multiBigwigSummary function in deeptools as described above for the ENCODE data comparison.

### BMMC dataset

#### Read mapping

Raw genomic reads were mapped and processed as described above for the cell culture single-cell dataset.

#### Quantification, quality control, and dimension reduction

Genomic bins were quantified using the AggregateTiles function in Signac, with binsize=5000 and min_counts=1 to quantify 5 kb bins genome-wide, retaining bins with at least one count. We retained cells with <10,000 and >100 H3K27me3 counts, and <10,000 and >75 H3K27ac counts for further analysis. We normalized the counts and reduced dimensionality for each assay by running the RunTFIDF, RunSVD, and RunUMAP functions in Signac and Seurat for each assay. We computed a WNN graph for H3K27me3 and H3K27ac using the FindMultiModalNeighbors function in Seurat, with reduction=list(“lsi.me3”, “lsi.ac”) and dims.list=list(2:50, 2:80) to use LSI dimensions 2 to 50 and 2 to 80 for H3K27me3 and H3K27ac, respectively. A 2-dimensional UMAP was created using the WNN graph by running the RunUMAP function in Seurat with nn.name=“weighted.nn” to use the pre-computed neighbor graph. We clustered cells using the WNN graph using the Smart Local Moving community detection algorithm (Waltman and van Eck 2013) (FindClusters function in Seurat with algorithm=3, resolution=3, graph.name=“wsnn”).

#### Cell annotation

To annotate cell types, we performed label transfer (Stuart et al. 2019) using the H3K27ac assay and a previously published scATAC-seq dataset containing healthy human bone marrow cells (Granja et al. 2019). As the original publication mapped reads to the hg19 genome, we re-processed the dataset from raw reads using the 10x Genomics cellranger-atac v2 software with default parameters, aligning to the hg38 genome. Code to reproduce this analysis is available on GitHub: https://github.com/timoast/MPAL-hg38. To transfer cell type labels from the scATAC-seq dataset to our multimodal NTT-seq dataset, we quantified scATAC-seq peaks using the H3K27ac assay, then performed TF-IDF normalization on the resulting count matrix using the IDF value from the scATAC-seq dataset. We performed LSI on the scATAC-seq BMMC dataset using the RunTFIDF and RunSVD functions in Signac with default parameters. We next ran the FindTransferAnchors function in Seurat, with reduction=“lsiproject”, dims=2:30, and reference.reduction=“lsi” to project the query data onto the reference scATAC-seq LSI using dimensions 2 to 30, and find anchors between the reference and query dataset. We ran TransferData with weight.reduction=bmmc_ntt[[“lsi.me3”]] dims=2:50 to weight anchors using LSI dimensions 2 to 50 from the H3K27me3 assay. We used these unsupervised cell type predictions as a guide when assigning cell clusters to cell types.

#### Trajectory analysis

We subsetted the BMMC dataset to contain cells annotated as HSPC, GMP/CMP, Pre-B, B, or Plasma cells. Using the subset object, we constructed a new UMAP dimension reduction by running FindTopFeatures, RunTFIDF, and RunSVD in Signac using the H3K27me3 assay. We converted the Seurat object containing these cells to a SingleCellExperiment object using the as.cell_data_set function in the SeuratWrappers package (https://github.com/satijalab/seurat-wrappers). We next ran Monocle 3 (Qiu et al. 2017) using the pre-computed UMAP dimension reduction by running the cluster_cells, learn_graph, and order_cells functions, setting the HSPC cells as the root of the trajectory. To find genomic features in each assay whose signal depended on pseudotime state, we quantified fragment counts for each cell in each 10 kb genome bin for the H3K27me3 and H3K27ac assays. To reduce the sparsity of the measured signal, we averaged counts for each genomic region across the cell’s 50 nearest neighbors, defined using the H3K27me3 neighbor graph with LSI dimensions 2 to 20, and normalized the fragment counts by the total neighbor-averaged counts per cell. For each genomic region we computed the Pearson correlation between the signal in the genomic region and the cell’s position in pseudotime. To find regions that underwent coordinated activation or repression we selected regions with an absolute difference in Pearson correlation between the H3K27me3 and H3K27ac assays greater than 0.5 (e.g., −0.25 correlation for H3K27me3 and +0.25 for H3K27ac). To display genomic regions in a heatmap representation we ordered cells based on their pseudotime rank and ordered genomic regions based on the position in pseudotime showing maximal H3K27me3 signal. For the purpose of visualization, we smoothed the signal for each genomic region by applying a rolling sum function with cells ordered based on pseudotime, summing the signal over 100-cell windows. This was performed using the roll_sum function in the RcppRoll R package (version 0.3.0).

We used the ClosestFeature function in Signac to identify the closest gene to each genomic region correlated with pseudotime. Genomic regions where the closest gene was >50,000 bp away were removed (21 genes for H3K27me3 and 7 genes for H3K27ac). To examine the gene expression patterns of these genes, we downloaded a previously integrated and annotated scRNA-seq dataset for the human bone marrow, produced as part of the HuBMAP consortium (https://zenodo.org/record/5521512) (Oetjen et al. 2018; Granja et al. 2019; HuBMAP Consortium 2019). We subset the scRNA-seq object to contain the same cell states that we examined in the NTT-seq data (HSC, LMPP, CLP, pro-B, pre-B, transitional B, naive B, mature B) and computed a gene module score for the active and repressed genes using the AddModuleScore function in Seurat.

## Acknowledgements

This work was supported by the National Institutes of Health (K99HG011489-01 to T.S.; RM1HG011014-01 to R.S., D.L.). B.Z. is a postdoctoral fellow of the Jane Coffin Childs Memorial Fund for Medical Research. This investigation has been aided by a grant from the Jane Coffin Childs Memorial Fund for Medical Research. We thank members of the Satija and NYGC Technology Innovation laboratories for feedback on the manuscript.

---

**Extended Data Figure 1.**
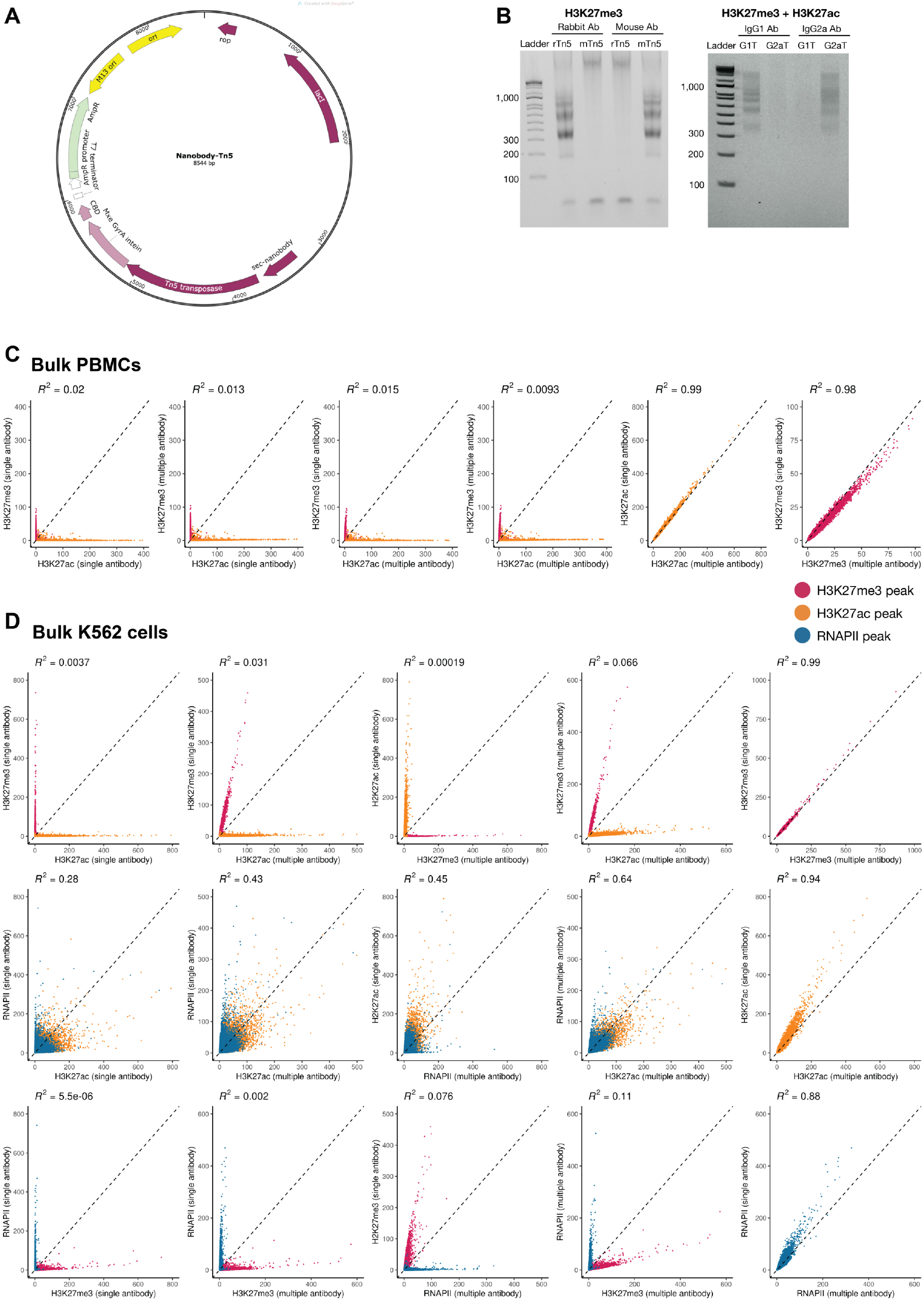
**A)** Nanobody-Tn5 fusion protein plasmid map schematic showing position of Tn5 and secondary nanobody sequences. **B)** Agarose DNA gel showing size-separation of PCR-amplified DNA sequencing library products for different combinations of nb-Tn5 and primary IgG antibody. Rabbit Ab: rabbit primary IgG antibody; Mouse Ab: mouse primary IgG antibody; IgG1 Ab: mouse IgG subtype 1 primary antibody; IgG2a Ab: mouse IgG subtype 2a primary antibody; rTn5: anti-rabbit IgG secondary nanobody-Tn5 fusion; mTn5: anti-mouse IgG secondary nanobody-Tn5 fusion; G1T: anti-mouse IgG1 secondary nanobody-Tn5 fusion; G2aT: anti-mouse IgG2a secondary nanobody-Tn5 fusion. Gels shows expected library amplification product (bands between 200 and 1,000 bp) in lanes where the nb-Tn5 fusion matches the primary IgG antibody (rabbit Ab + rTn5; mouse Ab + mTn5; IgG1 Ab + G1T; IgG2a Ab + G2aT). **C)** Scatterplots showing normalized fragment counts for H3K27me3 and H3K27ac peaks defined by ENCODE (ENCODE Project Consortium, 2012) for bulk multiplexed and non-multiplexed NTT-seq experiments in human PBMCs. Peaks are colored according to their chromatin modality (red: H3K27me3 peak, yellow: H3K27ac peak). Coefficient of determination (*R*2) between experiments are shown above each scatterplot. **D)** Scatterplots showing normalized fragment counts for H3K27me3, H3K27ac, and RNAPII peaks defined by ENCODE (ENCODE Project Consortium, 2012) for bulk multiplexed and non-multiplexed NTT-seq experiments in K562 cells. Peaks are colored according to their chromatin modality (red: H3K27me3; yellow: H3K27ac; blue: RNAPII).

**Extended Data Figure 2.**
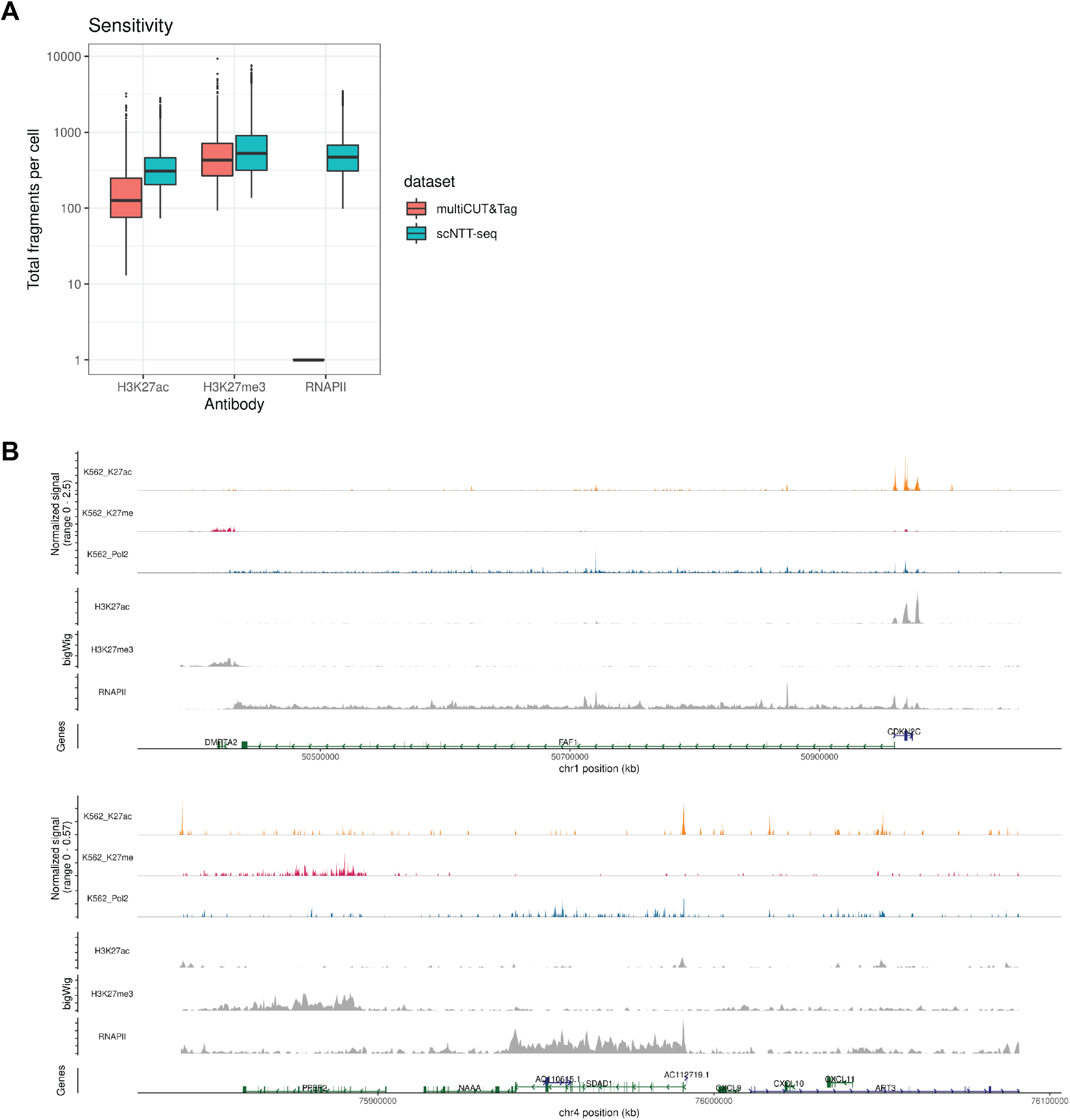
**A)** Total fragment counts per cell for multiCUT&Tag (Gopalan et al., 2021) and NTT-seq. Fragment counts on y-axis are on a log10 scale. multiCUT&Tag profiled only two marks, H3K27ac and H3K27me3, and so do not have RNAPII counts. **B)** Multimodal genome browser view of a representative genomic locus, for K562 cells. Top three tracks show H3K27ac, H3K27me3, and RNAPII profiled simultaneously in a single-cell experiment. Lower three tracks show H3K27ac, H3K27me3, and RNAPII profiled individually in bulk-cell NTT-seq experiments using K562 cells.

**Extended Data Figure 3.**
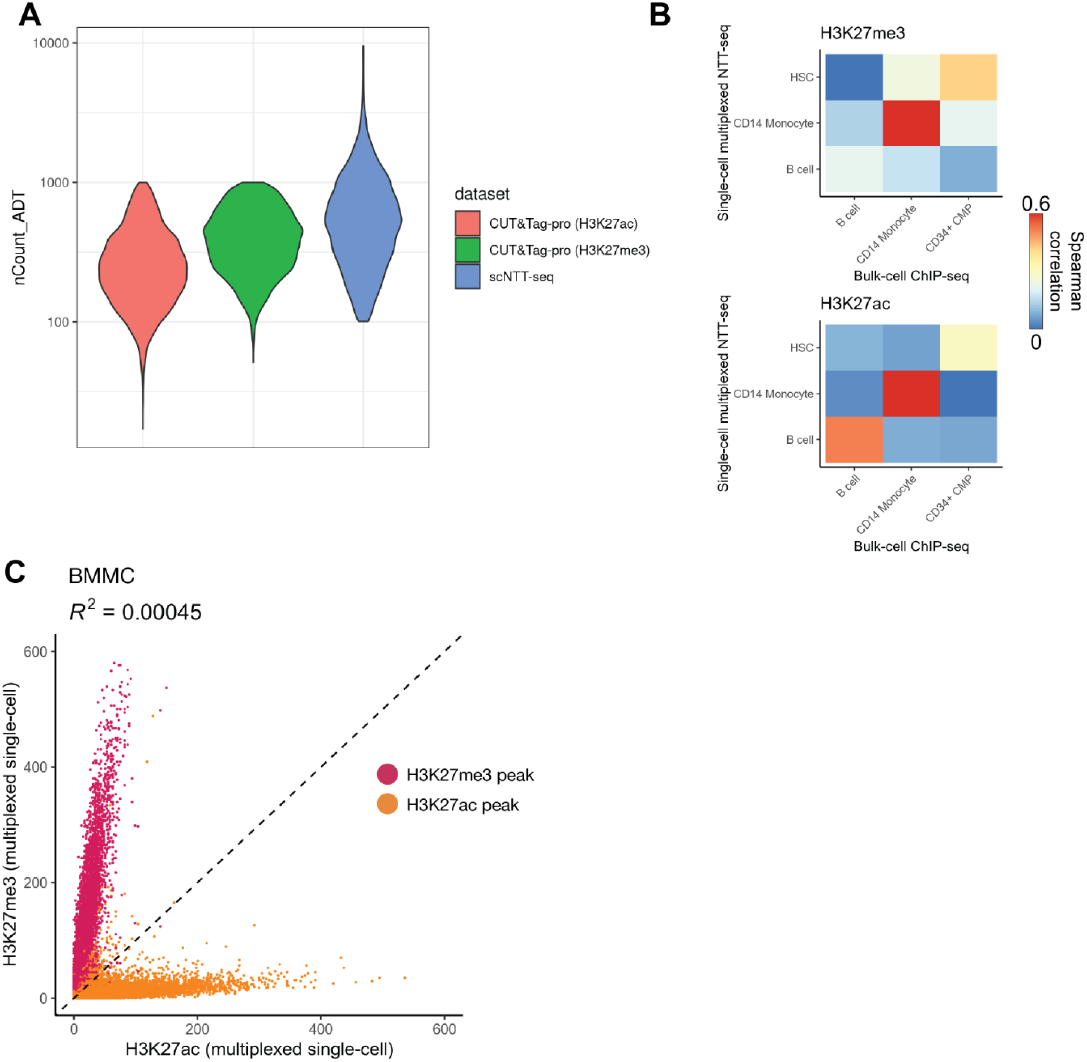
**A)** Comparison of total unique antibody-derived tag (ADT) counts sequenced per cell for CUT&Tag-pro (B. Zhang et al., 2021) and NTT-seq. **B)** Spearman correlation between H3K27me3 counts (top) or H3K27ac counts (bottom) for cells profiled using multiplexed single-cell NTT-seq, or FACS-sorted bulk ChIP-seq profiled by ENCODE (ENCODE Project Consortium, 2012). **C)** Scatterplot showing the number of counts per H3K27me3 and H3K27ac peak for each assay, for BMMC cells profiled using single-cell multiplexed NTT-seq. Peaks are colored according to their assay (red: H3K27me3 peaks; yellow: H3K27ac peaks).

## References

Amini, S. et al. (2014) ‘Haplotype-resolved whole-genome sequencing by contiguity-preserving transposition and combinatorial indexing’, Nature genetics, 46(12), pp. 1343– 1349. doi:10.1038/ng.3119.

Becht, E. et al. (2018) ‘Dimensionality reduction for visualizing single-cell data using UMAP’, Nature biotechnology. doi:10.1038/nbt.4314.

Carter, B. et al. (2019) ‘Mapping histone modifications in low cell number and single cells using antibody-guided chromatin tagmentation (ACT-seq)’, Nature communications, 10(1), p. 3747. doi:10.1038/s41467-019-11559-1.

Creyghton, M.P. et al. (2010) ‘Histone H3K27ac separates active from poised enhancers and predicts developmental state’, Proceedings of the National Academy of Sciences of the United States of America, 107(50), pp. 21931–21936. doi:10.1073/pnas.1016071107.

ENCODE Project Consortium (2012) ‘An integrated encyclopedia of DNA elements in the human genome’, Nature, 489(7414), pp. 57–74. Available at: http://www.nature.com/doifinder/10.1038/nature11247.

Gopalan, S. et al. (2021) ‘Simultaneous profiling of multiple chromatin proteins in the same cells’, Molecular cell, 81(22), pp. 4736–4746.e5. doi:10.1016/j.molcel.2021.09.019.

Granja, J.M. et al. (2019) ‘Single-cell multiomic analysis identifies regulatory programs in mixed-phenotype acute leukemia’, Nature biotechnology, 37(12), pp. 1458–1465. doi:10.1038/s41587-019-0332-7.

Hamilton, N.E. and Ferry, M. (2018) ‘ggtern: Ternary Diagrams Using ggplot2’, Journal of statistical software, 87, pp. 1–17. doi:10.18637/jss.v087.c03.

Hao, Y. et al. (2021) ‘Integrated analysis of multimodal single-cell data’, Cell, 184(13), pp. 3573–3587.e29. doi:10.1016/j.cell.2021.04.048.

Hassanzadeh-Ghassabeh, G. et al. (2013) ‘Nanobodies and their potential applications’, Nanomedicine, 8(6), pp. 1013– 1026. doi:10.2217/nnm.13.86.

Hawkins, J.A. et al. (2018) ‘Indel-correcting DNA barcodes for high-throughput sequencing’, Proceedings of the National Academy of Sciences of the United States of America, 115(27), pp. E6217–E6226. doi:10.1073/pnas.1802640115.

HuBMAP Consortium (2019) ‘The human body at cellular resolution: the NIH Human Biomolecular Atlas Program’, Nature, 574(7777), pp. 187–192. doi:10.1038/s41586-019-1629-x.

Janssen, S.M. and Lorincz, M.C. (2021) ‘Interplay between chromatin marks in development and disease’, Nature reviews. Genetics, pp. 1–17. doi:10.1038/s41576-021-00416-x.

Kaya-Okur, H.S. et al. (2019) ‘CUT&Tag for efficient epigenomic profiling of small samples and single cells’, Nature communications, 10(1), p. 1930. doi:10.1038/s41467-019-09982-5.

Kaya-Okur, H.S. et al. (2020) ‘Efficient low-cost chromatin profiling with CUT&Tag’, Nature protocols, 15(10), pp. 3264–3283. doi:10.1038/s41596-020-0373-x.

Li, H. et al. (2009) ‘The Sequence Alignment/Map format and SAMtools’, Bioinformatics, 25(16), pp. 2078–2079. doi:10.1093/bioinformatics/btp352.

Li, H. (2011) ‘Tabix: fast retrieval of sequence features from generic TAB-delimited files’, Bioinformatics, 27(5), pp. 718–719. doi:10.1093/bioinformatics/btq671.

Meers, M.P., Janssens, D.H. and Henikoff, S. (2021) ‘Multifactorial chromatin regulatory landscapes at single cell resolution’, bioRxiv. doi:10.1101/2021.07.08.451691.

Mimitou, E.P. et al. (2021) ‘Scalable, multimodal profiling of chromatin accessibility, gene expression and protein levels in single cells’, Nature biotechnology, pp. 1–13. doi:10.1038/s41587-021-00927-2.

Oetjen, K.A. et al. (2018) ‘Human bone marrow assessment by single-cell RNA sequencing, mass cytometry, and flow cytometry’, JCI insight, 3(23). doi:10.1172/jci.insight.124928.

Patro, R. et al. (2017) ‘Salmon provides fast and bias-aware quantification of transcript expression’, Nature methods, 14(4), pp. 417–419. doi:10.1038/nmeth.4197.

Picelli, S. et al. (2014) ‘Tn5 transposase and tagmentation procedures for massively scaled sequencing projects’, Genome research, 24(12), pp. 2033–2040. doi:10.1101/gr.177881.114.

Pleiner, T., Bates, M. and Görlich, D. (2018) ‘A toolbox of anti-mouse and anti-rabbit IgG secondary nanobodies’, The Journal of cell biology, 217(3), pp. 1143–1154. doi:10.1083/jcb.201709115.

Qiu, X. et al. (2017) ‘Reversed graph embedding resolves complex single-cell trajectories’, Nature methods, 14(10), pp. 979–982. Available at: http://www.nature.com/doifinder/10.1038/nmeth.4402.

Quinlan, A.R. and Hall, I.M. (2010) ‘BEDTools: a flexible suite of utilities for comparing genomic features’, Bioinformatics, 26(6), pp. 841–842. doi:10.1093/bioinformatics/btq033.

Ramirez, F. et al. (2014) ‘deepTools: a flexible platform for exploring deep-sequencing data’, Nucleic acids research, 42(W1), pp. W187–W191. Available at: https://academic.oup.com/nar/article-lookup/doi/10.1093/nar/gku365.

Saha, K., Bender, F. and Gizeli, E. (2003) ‘Comparative study of IgG binding to proteins G and A: nonequilibrium kinetic and binding constant determination with the acoustic waveguide device’, Analytical chemistry, 75(4), pp. 835–842. doi:10.1021/ac0204911.

Srivastava, A. et al. (2019) ‘Alevin efficiently estimates accurate gene abundances from dscRNA-seq data’, Genome biology, 20(1), p. 65. doi:10.1186/s13059-019-1670-y.

Stuart, T. et al. (2019) ‘Comprehensive Integration of Single-Cell Data’, Cell, 177(7), pp. 1888–1902.e21. doi:10.1016/j.cell.2019.05.031.

Stuart, T. et al. (2021) ‘Single-cell chromatin state analysis with Signac’, Nature methods, pp. 1–9. doi:10.1038/s41592-021-01282-5.

Tedesco, M. et al. (2021) ‘Chromatin Velocity reveals epigenetic dynamics by single-cell profiling of heterochromatin and euchromatin’, Nature biotechnology, pp. 1–10. doi:10.1038/s41587-021-01031-1.

Tie, F. et al. (2009) ‘CBP-mediated acetylation of histone H3 lysine 27 antagonizes Drosophila Polycomb silencing’, Development, 136(18), pp. 3131–3141. doi:10.1242/dev.037127.

Vasimuddin, M. et al. (2019) ‘Efficient Architecture-Aware Acceleration of BWA-MEM for Multicore Systems’, in 2019 IEEE International Parallel and Distributed Processing Symposium (IPDPS). ieeexplore.ieee.org, pp. 314–324. doi:10.1109/IPDPS.2019.00041.

Waltman, L. and van Eck, N.J. (2013) ‘A smart local moving algorithm for large-scale modularity-based community detection’, The European physical journal. B, 86(11), p. 471. doi:10.1140/epjb/e2013-40829-0.

Wang, Q. et al. (2019) ‘CoBATCH for High-Throughput Single-Cell Epigenomic Profiling’, Molecular cell [Preprint]. doi:10.1016/j.molcel.2019.07.015.

Zaborowska, J., Egloff, S. and Murphy, S. (2016) ‘The pol II CTD: new twists in the tail’, Nature structural & molecular biology, 23(9), pp. 771–777. doi:10.1038/nsmb.3285.

Zhang, B. et al. (2021) ‘Characterizing cellular heterogeneity in chromatin state with scCUT&Tag-pro’, bioRxiv. doi:10.1101/2021.09.13.460120.

Zhang, H. et al. (2021) ‘Fast alignment and preprocessing of chromatin profiles with Chromap’, Nature communications, 12(1), p. 6566. doi:10.1038/s41467-021-26865-w.

Zhang, Y. et al. (2008) ‘Model-based analysis of ChIP-Seq (MACS)’, Genome biology, 9(9), p. R137. Available at: http://genomebiology.biomedcentral.com/articles/10.1186/gb-2008-9-9-r137.13

